# On the identifiability of transmission dynamic models for infectious diseases

**DOI:** 10.1101/021972

**Authors:** Jarno Lintusaari, Michael U. Gutmann, Samuel Kaski, Jukka Corander

## Abstract

Understanding the transmission dynamics of infectious diseases is important for both biological research and public health applications. It has been widely demonstrated that statistical modeling provides a firm basis for inferring relevant epidemiological quantities from incidence and molecular data. However, the complexity of transmission dynamic models causes two challenges: Firstly, the likelihood function of the models is generally not computable and computationally intensive simulation-based inference methods need to be employed. Secondly, the model may not be fully identifiable from the available data. While the first difficulty can be tackled by computational and algorithmic advances, the second obstacle is more fundamental. Identifiability issues may lead to inferences which are more driven by the prior assumptions than the data themselves. We here consider a popular and relatively simple, yet analytically intractable model for the spread of tuberculosis based on classical IS6110 fingerprinting data. We report on the identifiability of the model, presenting also some methodological advances regarding the inference. Using likelihood approximations, it is shown that the reproductive value cannot be identified from the data available and that the posterior distributions obtained in previous work have likely been substantially dominated by the assumed prior distribution. Further, we show that the inferences are influenced by the assumed infectious population size which has generally been kept fixed in previous work. We demonstrate that the infectious population size can be inferred if the remaining epidemiological parameters are already known with sufficient precision.

---

November 19, 2015

Statistical models for transmission dynamics are widely employed to answer fundamental questions about infectivity of bacteria and viruses, and to make predictions for intervention policies, such as vaccines, de-colonization and case containment. For some types of infectious disease, the complexity of the transmission process and the corresponding model, combined with the characteristics of the available data, make the inference an intricate task. A particular difficulty arises from the need to use computationally intensive methods. Examples include the work by Tanaka *et al*. 2006; Sisson *et al*. 2007; Blum 2010; Stadler 2011; Fearnhead and Prangle 2012; Del Moral *et al*. 2012; Baragatti *et al*. 2013; Albert *et al*. 2015, who considered the transmission dynamics of *Mycobacterium tuberculosis* based on IS6110 fingerprinting data from tuberculosis (*M*. *tuberculosis*) cases in San Francisco, reported earlier by Small *et al*. (1994). Except for Stadler (2011), who proposed an inference scheme based on likelihood and Markov chain Monte Carlo, the above-mentioned studies employed and improved an approximate inference technique known as approximate Bayesian computation (ABC), which was originally introduced by Tavaré *et al*. (1997).

Although the estimation of epidemiological parameters of *M. tuberculosis* with the model of Tanaka *et al*. (2006) has been widely studied, concerns with identifiability have been raised. Originally Tanaka *et al*. (2006) reported a wide credible interval for the reproductive value *R* and later, Blum (2010) suggested that the data of Small *et al*. (1994) are not informative enough for confident estimation of *R* in the original setting by visually comparing the prior to the inferred posterior distribution. Stadler (2011) further questioned the accuracy of the ABC approach of Tanaka *et al*. (2006) after obtaining significantly different estimates with her method. This concern was later reconciled by Aandahl *et al*. (2014) who showed that the ABC method was valid and, moreover, also computationally more efficient. However, their confirmatory experiments with synthetic data were only covering the setting with a single free parameter, which leaves the question about estimability open for models with multiple free parameters.

In certain cases the outcomes of ABC may not be accurate because the method includes several approximations and because practical algorithms can have several tuning parameters. Multiple validation methods have thus been developed. Many operate by using synthetic data generated with known parameter values in place of the observed data, and comparing the inference results with the known parameter values. Wegmann *et al*. (2009), for example, used the absolute difference between the data-generating parameter values and the posterior point estimates to test whether the inferred posterior distribution is concentrated around the right parameter values; to test whether the spread of the distribution is not overly large or small, they suggest to compute the proportion of times the credible interval contain the data-generating parameter values (see also the work by Prangle *et al*. (2014)). Csilléry *et al*. (2012), on the other hand, recommend comparing the observed data with data simulated from the posterior predictive distribution. Further validation methods include confidence intervals, interquantile ranges, visualizations of the posterior distributions (e.g. Tanaka *et al*. 2006; Toni *et al*. 2009; Blum 2010), and principal component analysis (Toni *et al*. (2009) and Cornuet *et al*. (2010)).

Failure to pass some of the validation tests can be due to several reasons. The approximation may not be accurate enough, the settings of the inference algorithm could be the problem, or the issue could be deeper: the model may not be fully identifiable in the first place. In this paper, we assess the identifiability of the epidemiological model of Tanaka *et al*. (2006) for genotype data of the kind available from the San Francisco study (Small *et al*. 1994). Since the likelihood function indicates the informativeness of the data, we approach the identifiability problem directly by approximating the likelihood. Further, as previous ABC studies dealing with the same epidemiological model have assumed a fixed infectious population size for the data, we investigate how this choice influences the estimation of the epidemiological parameters, and whether it is possible to infer the population size from this kind of genotype data without access to more extensive surveillance data about incidence. Since comparable data are widely considered for many different kinds of pathogens, the issue of model identifiability – and our approach of addressing it by approximating the likelihood function – is of wider general interest beyond the particular case discussed here.

## MODELS AND METHODS

**Model for disease transmission:** The model considered in the paper is a linear birth-death process with mutations (BDM) introduced by Tanaka *et al*. (2006). The process model is defined as follows: Each infected individual, hereafter host, carries the pathogen characterized by an allele at a single locus of its genome. The host transmits the pathogen and the corresponding allele with rate *α*, and dies or recovers with rate *δ*. For simplicity we call *α* the birth rate and *δ* the death rate. In addition, the pathogen mutates within the host with rate *τ*, resulting each time in a novel allele in the population of hosts (infinite alleles model). When simulating the process, one begins with a single host and stops when either the population of hosts *X* reaches a predetermined size *m* or the pathogen goes extinct. The observation model assumes sampling of *n <* m hosts from *X* without replacement. It has been earlier noted by Stadler (2011) that due to the time scaling of the model, at least one of the rate parameters must be fixed. Similarly to many of the earlier studies, we use a time scale of one year and fix the mutation rate to *τ* = 0.198 per year throughout the experiments. Likewise, the infectious population size *m* is set to 10, 000 unless otherwise stated.

The epidemiological parameters of interest in this study are the reproductive value *R* and the net transmission rate *t*. In addition, we will consider inference of the underlying infectious population size *m* given some estimate of *R* and *t*. In what follows, we will often use *θ* to denote the tuple (*R, t*). The epidemiological parameters *R* and *t* are in a one-to-one correspondence to the event rate parameters of the BDM process: *R* = *α/δ, t* = *α − δ,* and *δ* = *t/*(*R−*1), *α* = *tR/*(*R−*1).

**Data:** The alleles of the pathogen carried by the n sampled hosts are summarized in the form of the allele vector 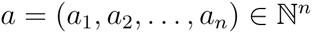, where element *a_i_* equals to the number of allele clusters of size *i* present in the sample. An allele cluster is a set of hosts having the same allele of the pathogen, and its size is the number of hosts which belong to the cluster. For example, the vector *a* = (4, 0, 1) implies that there are four singleton clusters and one cluster with three hosts in the sample. In other words, there are four different alleles each found in only one host and one allele shared by three hosts. The size of *a* is defined as the sample size *n*, which can be written in terms of *a* as *n* = Σ*_i_ ia_i_.*

For inference of the parameters, as in Tanaka *et al*. (2006), we used the San Francisco data of Small *et al*. (1994) which consist of an allele vector *a\** of size *n* = 473. Its nonzero elements are 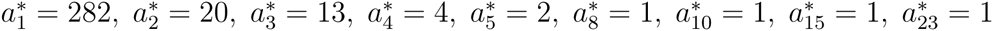, and 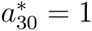

**Inference method:** The likelihood function plays a central role in statistical inference. But for the model considered in this paper, it cannot be expressed analytically in closed form (Tanaka *et al*. 2006). Tanaka *et al*. (2006) thus used approximate Bayesian computation (ABC) for the inference. We here approximate the likelihood function using kernel density estimation with a uniform kernel and a distance measure *d*, an approach which is related to ABC but which makes explicit its inherent approximations (Blum 2010; Gutmann and Corander 2015).

For a fixed value of the population size *m*, the likelihood function *L*(*θ*) is approximated as 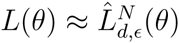,

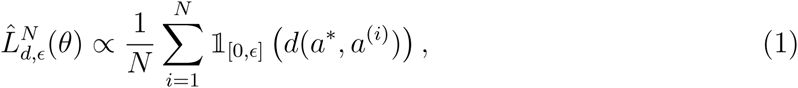
where 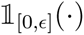 is an indicator function which equals one if the distance *d* is less than a threshold *ϵ* and zero otherwise, *a*^(^*^i^*^)^ is an allele vector simulated using parameter *θ*, and *N* is the number of such simulations performed (Gutmann and Corander 2015). The distance *d*(*a*, a*) is a non-negative function which measures the similarity between the observed allele vector *a*\* and the simulated allele vector *a.* Possible choices of *d* are discussed below. Equation 1 means that the likelihood is approximated by the fraction of times the simulated allele vector *a* is within distance *ϵ* from the observed allele vector *a*\*. The approximate likelihood function for inference of the population size *m* for fixed *θ* is defined in an analogous manner.

While there are several variants of the inference procedure of ABC, they are essentially built out of sampling candidate parameter values *θ* and retaining those for which the distance *d*(*a*,a*) is less than the threshold *ϵ*. Under certain conditions, the retained parameters correspond to samples from the posterior. In ABC, the approximate likelihood function is generally never explicitly constructed, but is implicitly represented by the sampled parameter values for which the simulated data are close to the observed data. By contrast, in this paper the likelihood function is explicitly approximated using Equation 1, which is important because it provides information about the identifiability of the parameters.

Since the parameter space is low-dimensional, we can evaluate the approximate likelihood function by varying the parameters on a grid. Several grids were created based on the different inference tasks considered.

- For inference of *θ*, we formed a 137 × 120 evenly spaced grid over a subspace Δ*_a_* × Δ*_δ_* = [0.3, 2] × [0.0125,1.5] of the (*α, δ*) BDM parameter space. At each node of the grid, *N* = 3000 allele vectors were simulated in order to approximate the likelihood in that location by using Equation 1.
- To study the effect of *m* on *R*, four different grids (one for each value of *m*) with 41 nodes were created, being subsets of the interval Δ*_α_* = [0.53, 1.4] and having *N* = 3000 simulations in each node.
- For the evaluation of the feasibility of inference of *m* also four grids were used, two with size 50 and two with 51, *N* = 3000, all grids being subsets of the interval Δ*^N^* = [500, 25000].
- For the final inference of the population size *m*, we used two grids, one with 15 nodes and the other with 26 nodes, both subsets of Δ*^N^* = [500, 35000] with *N* = 30, 000 simulated allele vectors in each node.

The amount of simulated data in the grids above was selected to ensure the stability of the likelihood approximations. Results for the stabilization of the marginal likelihoods for the grid over the (*α, δ*) parameter space above are shown in Appendix A. The dimensions of the two dimensional grid were chosen such that the grid includes the modes of the likelihoods used in the experiments. The one dimensional grids were chosen so that they included all of the significant mass of the approximated likelihood functions, so that numerical computation of the posterior means is possible.

**Distance measures used to approximate the likelihood:** The likelihood approximation in Equation 1 relies on a distance measure *d*(*a*, a*) between the observed sample *a*\* and the sample *a* produced by the simulation process with the parameter vector *θ*. Consequently, different distance measures may lead to different estimation results, depending on how much information about the generating process they are able to capture from the data. It is thus natural to ask which distance measures would be optimal for a model of the type considered here.

In the limit of *ϵ* → 0 and *N → ∞,* one can easily define distance measures *d*(*a*, a*) which lead to exact likelihoods *L*(*θ*). The only requirement is that *d*(*a*, a*) = 0 if and only if *a* = *a*.* In practice, however, too small thresholds *ϵ* and a very large number of simulations *N* are computationally not feasible. Therefore one has to rely on likelihood approximations 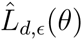 dictated by the distance measure *d*, a non-zero threshold value *ϵ* and a finite *N*.

Given a fixed number of simulations *N*, differences in the quality of the approximations arise from the ability of the distance measures *d* to produce approximate likelihood functions which are as close as possible to L(*θ*). Because the likelihood L(*θ*) is unknown in the first place, the evaluation of the approximations is challenging. One method to evaluate the goodness of an approximate likelihood function is to measure the goodness of the corresponding estimates using synthetic observed data *a^s^* where the data generating parameters *θ^s^* are known. In particular, the mode of the likelihood approximation should, on average, be near *θ^s^* if the sample *a^s^* is in general informative enough.

In this study, we evaluate the performance of three different distance measures. The baseline distance measure is the one introduced by Tanaka *et al*. (2006) and is defined as

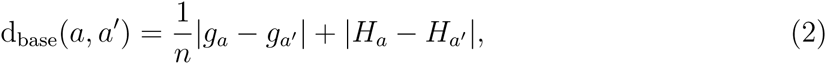
where *H_a_* = 1 − Σ*_i_ a_i_*(*i/n*)^2^ is a gene diversity summary statistic and *g_a_* = Σ*_i_ a_i_* is the number of distinct alleles in *a*, or in other words the total number of clusters present in the data.

The second distance measure called “simple” is defined as

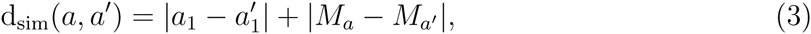
where *a* and *a′* are allele vectors and *M_a_* = max {*i*|*a_i_* ≠ 0} is the largest cluster size in *a*. This measure compares the number of singleton clusters in the sample and the sizes of the largest clusters. It can be seen as a simplified version of the baseline distance measure by excluding some information.

The third distance measure d^GKL^ is motivated by the observation that element *a_i_* of the allele vector *a* is proportional to the frequency of occurrence of cluster size *i* in the data, with the proportionality factor *g_a_* = Σ*_i_ a_i_* being equal to the total number of clusters present. We can thus consider *a_i_* to indicate the probability to observe cluster size *i* in the data. This probabilistic interpretation of vector *a* opens up the possibility to use known divergence measures from probability theory to gauge the similarity between two vectors *a* and *a′*. In more detail, since both *a* and *a*′ correspond to estimated probabilities, we shall discount small differences in the number of clusters of similar size, which may be present due to chance alone, and represent *a* and *a*′ by smooth continuous functions *f_a_* and *f_a'_* approximating the two vectors in the discrete locations *i*, that is, *f_a_*(*i*) ≈ *a_i_* and 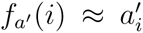. Like *a* and *a′* correspond to unnormalized probability vectors, *f_a_* and *f_a'_* correspond to unnormalized probability densities, and we can assess their similarity by the generalized Kullback-Leibler divergence,

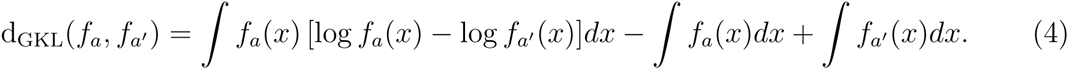

The generalized Kullback-Leibler divergence belongs to the family of Bregman divergences (Bregman 1967) which have a number of desirable properties, for example nonnegativity and being equal to zero if *f_a_* = *f_a'_*, as well as useful applications (see for example Collins *et al*. 2002; Frigyik *et al*. 2008; Gutmann and Hirayama 2011). Unlike in the usual Kullback-Leibler divergence, the integrals over *f_a_* and *f_a'_* need not equal one in the generalized Kullback-Leibler divergence. In fact, by construction of *f_a_* and *f_a'_*, the difference between their integrals assesses the difference between the total number of clusters *g_a_* and *g_a'_*, much like in d_base_. Conceptually, our probabilistic interpretation of *a* boils down to using the cluster size as a summary statistic and comparing its probability distribution for observed and simulated data in a nonparametric way.

## RESULTS

**Evaluations of the distance measures:** In order to compare the performance of the alternative distance measures d_sim_ and d^GKL^ to the baseline distance measure d_base_ in likelihood approximation, the difference Δ_error_ between the relative errors in their respective estimates was computed. A value Δ_error_ > 0 indicates that the relative error is larger for the baseline compared with the alternative method, in which case the alternative method would be preferable.

The relative error was defined as 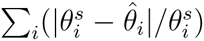, where *θ*^s^ is the true data-generating parameter value used to simulate the synthetic data *a^s^,* and 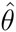 is the estimate obtained by maximizing the approximate likelihood. Maximization was performed in a simple way, by searching for the maximal value over the two-dimensional grid. To compute the relative error, a total of 50 synthetic observations *a^s^* were simulated using *θ*^s^, and the likelihood function was approximated with the grid for each observation *a^s^.* The threshold *ϵ* was set for each distance measure to the value minimizing the sum of the relative errors over the 50 trials.

We considered three different setups for the data-generating parameter value *θ*^s^. In the first setup, the value of *θ^s^* was set to the estimate (3.4, 0.69) of Tanaka *et al*. (2006). The second setup uses the estimate (2.1, 0.57) from Aandahl *et al*. (2014). To further see if the values of the actual epidemiological parameters had an effect on the estimation accuracy, we considered one more setup where both the reproductive value *R* and the transmission rate *t* of the first setup were divided by two.

Figure 1 shows the resulting distribution of Δ_error_ for the comparison between d_sim_ and d_base_ (blue curve), and the comparison between d^GKL^ and d_base_ (red curve). The simple distance measure performs slightly worse than the baseline although the difference is not significant in any of the setups (the null hypothesis of a zero mean of Δ_error_ cannot be rejected). This means that reducing the distance measure d_base_ of Tanaka *et al*. (2006) to the simpler d_sim_ does not cause a notable reduction in estimation accuracy. The generalized Kullback-Leibler distance d^GKL^, on the other hand, performs slightly better than the baseline and the difference is significant in the last setup (the zero mean hypothesis of Δ_error_ can be rejected at the *p*-value 0.0237). It should nevertheless be noted that the absolute errors tend to be rather large with all of the distance measures as shown in Appendix B.

**Figure 1:**
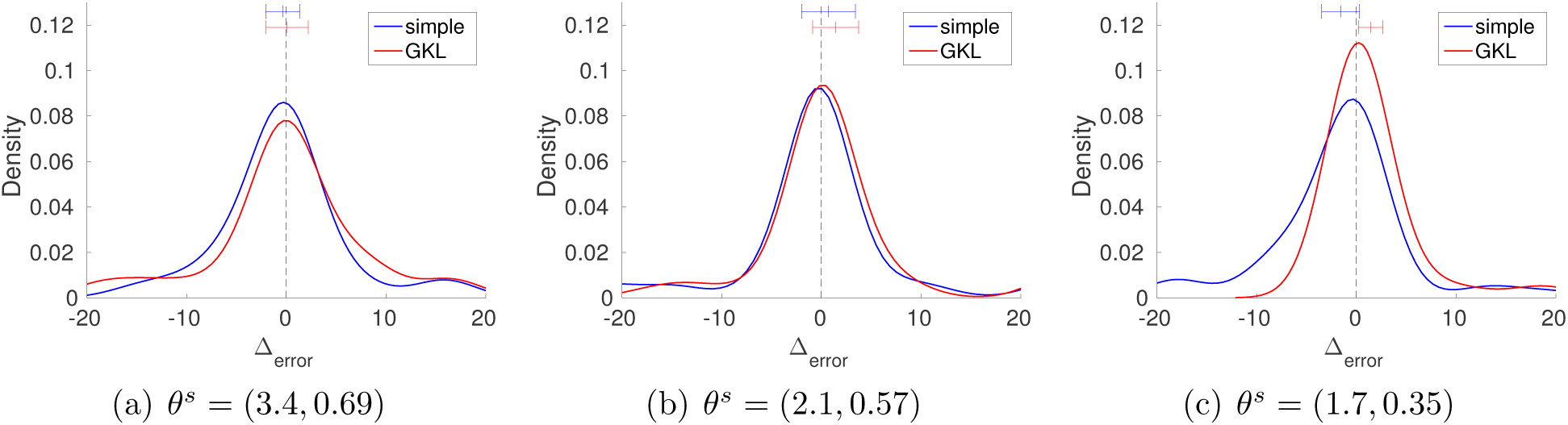
The distribution of the difference Δ_error_ between the relative estimation errors for the baseline and two alternative distance measures. A positive value of Δ_error_ indicates that the alternative method performs better. The intervals at the top show the estimated mean and the 95% confidence interval.

Since d^GKL^ was found to perform at least as well as the other measures, it was used in the remaining parts of the paper unless stated otherwise. Furthermore, we will for simplicity often drop the qualifier “approximate” and use “approximate likelihood” and “likelihood” interchangeably.

**Relative effects of likelihood and prior on the posterior:** The simulator operates genuinely in the (*α, δ*) space, where *α* and *δ* are the birth rate and the death rate in the model. Accordingly, all of the ABC studies so far have assumed an uninformative (uniform) prior for the region 0 *< δ < α* in the Bayesian framework. The law of transformation of random variables implies that choosing a uniform prior for (*α, δ*) is equivalent to choosing the following prior for the epidemiological parameters (R, t),

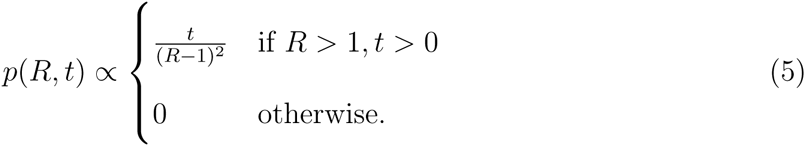

The prior is visualized in Appendix C. The formula and the figure show that its probability mass is concentrated on small values of the reproductive value *R*.

Figure 2(a) shows the likelihood function of (*R, t*) for the San Francisco data of Small *et al*. (1994) on the rectangle defined by 1.2 *< R <* 80 and 0.4 *< t <* 0.7. We used the same grid as in the previous section with threshold equal to the 10^−4^ quantile of all the distances. The likelihood function is flat over large areas of the parameter space and many values of *R* are equally likely, which means that the data are not informative enough to identify the parameter *R* of the model.

**Figure 2:**
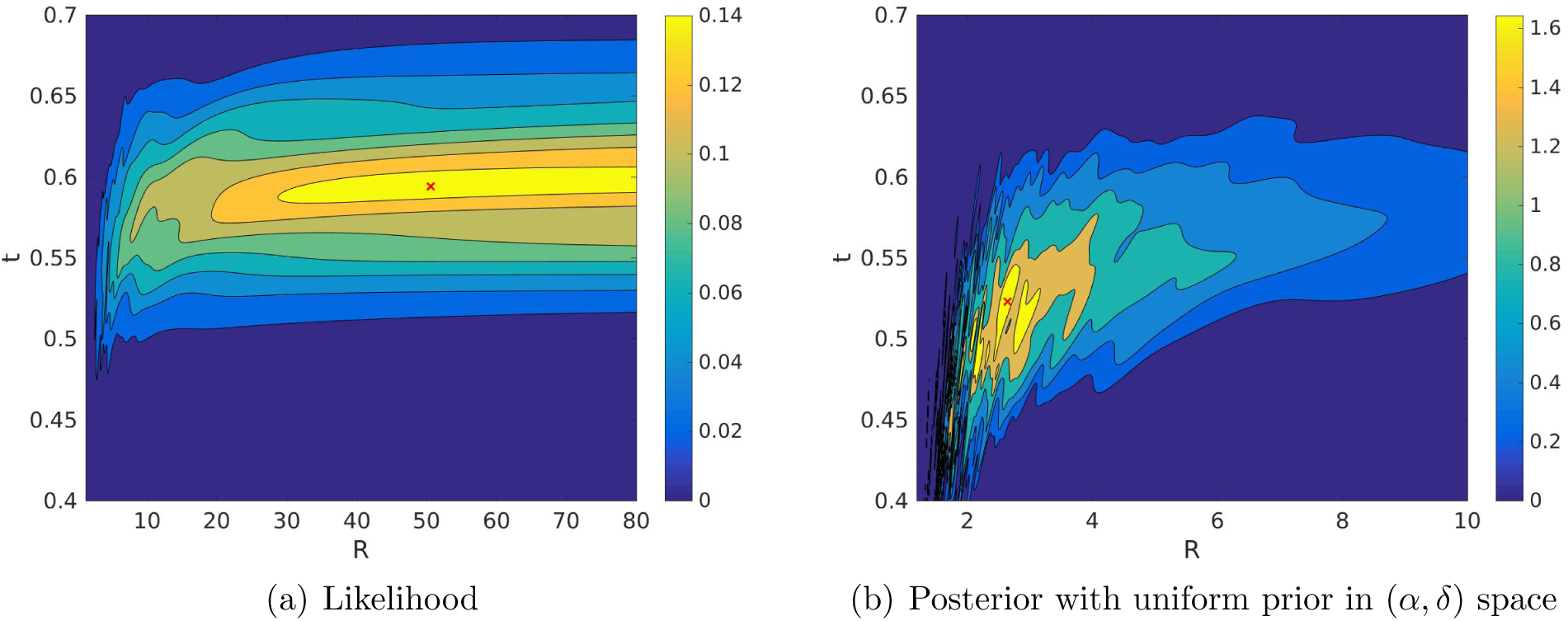
Likelihood and posterior distribution when using a uniform prior in the (*α, δ*) space. The approximate posterior is obtained by multiplying the approximate likelihood with the prior on the grid. The red cross denotes the mode. Note the different scales of the x-axes.

Figure 2(b) shows the posterior distribution of (*R, t*) for the prior in Equation 5, that is, for the uniform prior on (*α, δ*). It can be seen that the prior leads to a substantial shift of the probability mass towards the lower end of values of *R*. The difference between the modes of the likelihood and the posterior is striking: *R* = 50.6 versus *R* = 2.7 as shown in Table 1. The table also shows the posterior means and credible intervals for the case that the likelihood is interpreted as the posterior distribution with a uniform prior in the (*R, t*) space. It should be noted that the upper value of the credible intervals for *R* is an artifact of limiting our computation to values of *R* less than 80. The shape of the likelihood in Figure 2 (a) suggests that computations with larger values of *R* would lead to a corresponding increase of the credibility intervals. The results mean that the prior dominates the posterior distribution, which confirms previous findings by Blum (2010).

**Table 1:**
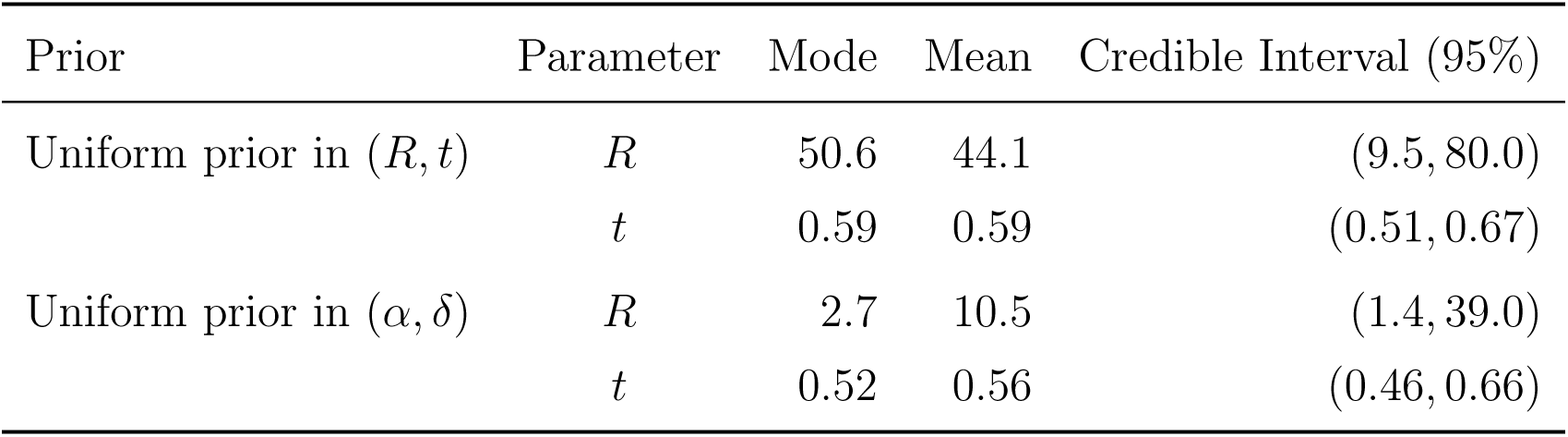
Effect of the prior on the posterior mode, mean, and credible interval of the epidemiological parameters of *M. tuberculosis* for the San Francisco data.

**Effect of the infectious population size:** The previous sections suggest that data of the kind considered in the San Francisco study do not carry enough information for accurate inference of *R*, but that the prior plays a major role. To make the inference of *R* possible, Aandahl *et al*. (2014) obtained an estimate 0.52 for the death rate *δ*by summing the rates of self cure, death from causes other than tuberculosis and death from untreated tuberculosis as estimated in other studies. The parameter *δ* was then either fixed to this value or an informative prior centered to it was used. Also previously, in all of the corresponding ABC studies, the infectious population size *m* has been fixed, commonly to the value *m* = 10, 000 following Tanaka *et al*. (2006). We were thus interested in whether reducing the infectious population size to a smaller, possibly more realistic number has an influence on the estimated value of *R*. To ease the comparison with previous studies, we used the distance dbase in the likelihood approximation. The thresholds were set to the 5 · 10^−3^ quantile of the distances in the respective grids.

Figure 3 shows the likelihoods of *R* for *m* ∈ {1000, 5000, 10000, 20000} using the San Francisco data (Small *et al*. 1994) and *δ* = 0.52. The difference in location of the likelihoods is clear with modes shifting to the right with increasing *m*. The mode locations were (1.1, 1.6, 1.9, 2.2) given in the order of increasing *m*. Posterior means with uniform prior over the support of the likelihoods were the same. The results show that the assumed infectious population size *m* affects the inference of *R*.

**Figure 3:**
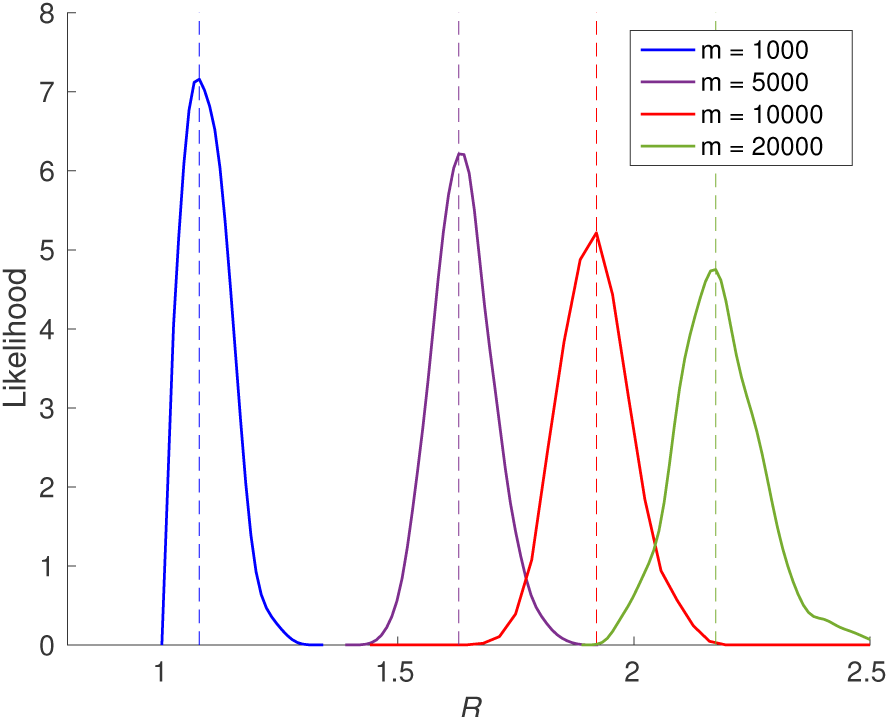
Likelihoods of the reproductive value *R* with fixed death rate *δ* = 0.52 and four alternative population sizes *m* using the San Francisco data. The vertical lines indicate the modes of the likelihoods.

**Inference of the infectious population size:** The observed effect of the infectious population size *m* on the inference of *R* means that there is some (statistical) dependency between *m* and *R*. This suggests that it might be possible to infer the size of the underlying infectious population from the data when *R* and *δ* are known. Alternatively, due to the relationship between the parameters, knowing any two of *α, δ*, *R* or *t* would be sufficient.

We fixed *δ* = 0.52 as earlier and considered two alternative configurations: *R* = 2.1 and *R* = 1.1. The former is also the estimate of Aandahl *et al*. (2014). To test whether inference of the infectious population size parameter is possible, we ran 50 trials with synthetic data as before, and determined each time the maximizer of the approximate likelihood on the grid. The thresholds *ϵ* were set to the 10^−2^ quantile of the distances. Table 2 shows the results of these experiments: The estimated population sizes are reasonable, and the actual population size *m* is covered by the 95% confidence interval of the mean in all but one of the cases. Only in the last case, the true *m* is just barely outside of the interval. These results thus illustrate that estimation of *m* is possible, provided that reliable information is available about the other epidemiological parameters.

**Table 2:** Mean estimated population size 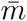 for 50 trials and the respective confidence interval (CI) under two alternative configurations of *R* and population size *m*. Death rate was fixed to *δ* = 0.52. Results are for synthetic data.

| | $m$ | $\bar{m}$ | CI (95%) |
| --- | --- | --- | --- |
| $R = 2.1$ | 1000 | 1051 | (990, 1112) |
|  | 10000 | 10020 | (9380, 10660) |
| $R = 1.1$ | 1000 | 1007 | (976, 1038) |
|  | 10000 | 10510 | (10003, 11017) |

We next estimated the infectious population size for the *M. tuberculosis* in San Francisco area during the time the data of Small *et al*. (1994) were collected. Threshold *ϵ* was selected as the 10^−3^ quantile of the distances. Figure 4 shows the likelihood functions for two different values of *R*: *R* = 1.1 and *R* = 2.1 (with *δ* = 0.52 as before). Assuming *R* = 1.1 produced the posterior mean 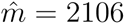 and the 95% credible interval (1166, 3226) with uniform prior over the support of the likelihood function. Assuming *R* = 2.1, on the other hand, produced the posterior mean 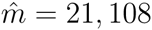 and the 95% credible interval (13408, 30155).

**Figure 4:**
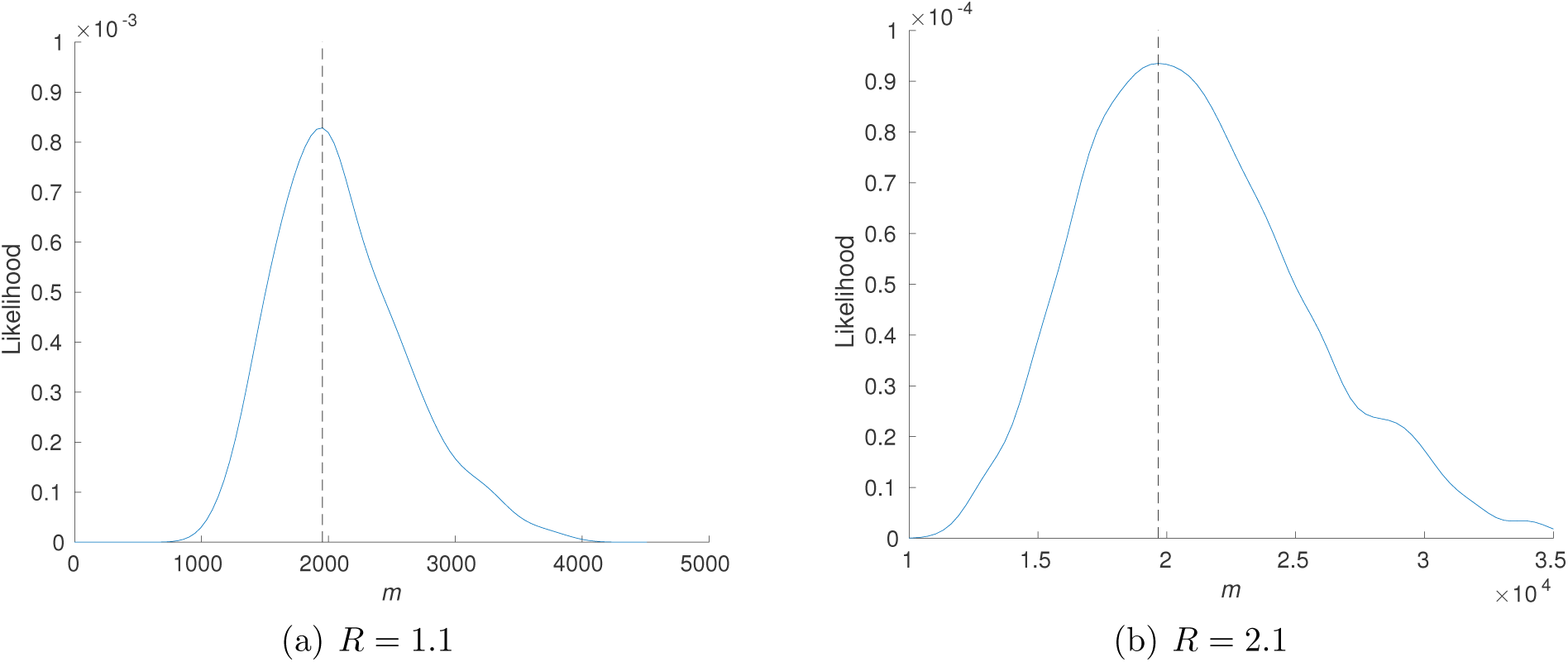
Likelihoods of population size *m* with fixed death rate *δ* = 0.52 and two alternative values of *R* using the San Francisco data. The vertical lines indicate the modes of the likelihoods.

**Figure 5:**
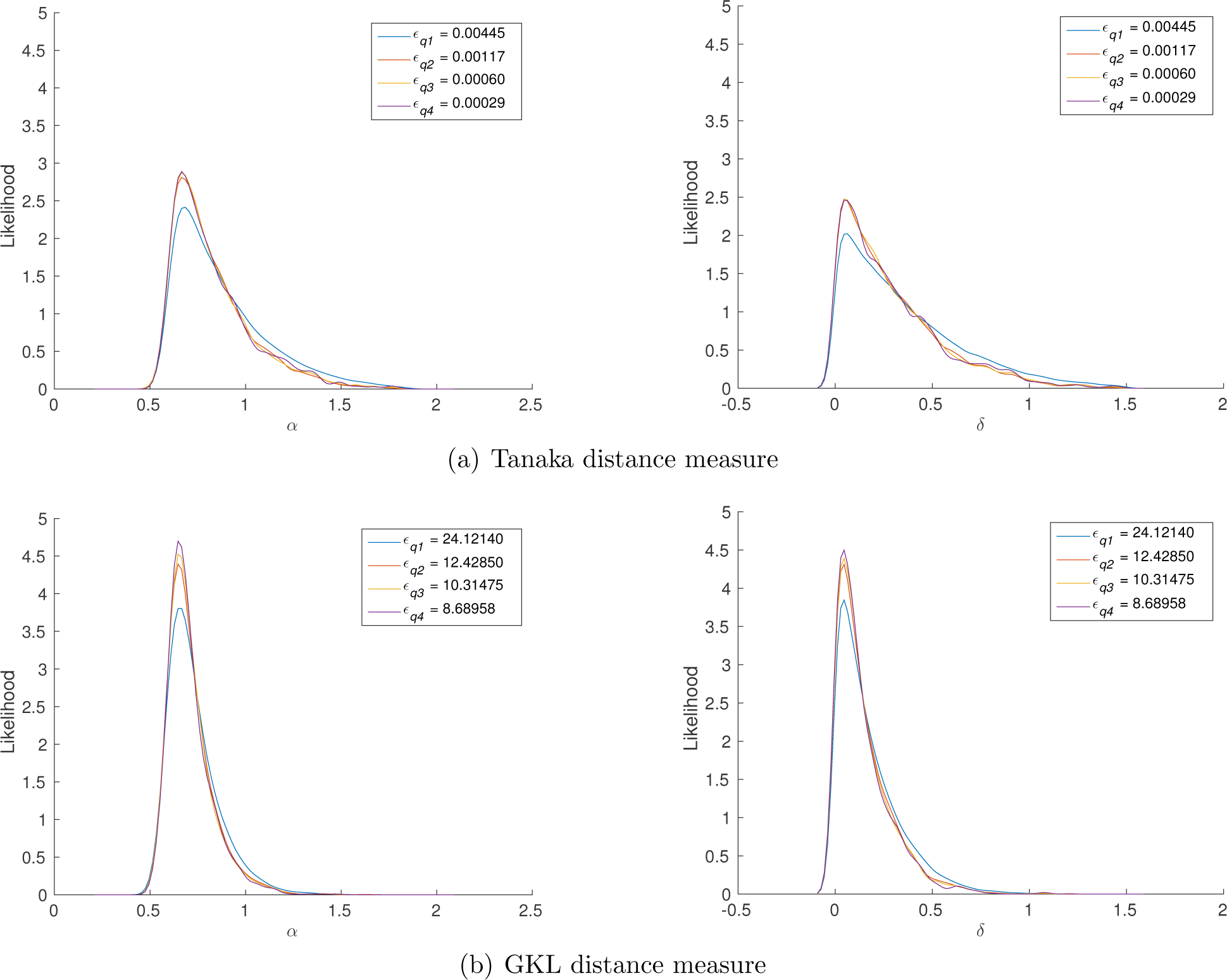
Stabilization (convergence) of the approximate marginal likelihoods for decreasing thresholds. The thresholds *ϵ_qi_* were obtained from the quantiles (*q*_1_, *q*_2_, *q*_3_, *q*_4_) = (0.001, 0.0001, 0.00005, 0.000025) of the distribution of the distances.

## DISCUSSION

Statistical inference plays an important role in the study of the transmission dynamics of infectious diseases. In this paper, we considered some of the challenges arising from model identifiability and from the expert choices necessary for approximate inference, using the relatively simple, yet analytically intractable model of Tanaka *et al*. (2006) for the transmission dynamics of *M. tuberculosis*. It is reasonable to assume that these problems persist for more complex transmission models, unless molecular and epidemiological data are detailed enough to mitigate their effect.

Due to the intractability of the transmission model, an approximate inference approach was used, belonging to the framework of “approximate Bayesian computation” (ABC), which relies on a distance measure gauging similarity between observed and simulated data. An alternative approach has been presented by Stadler (2011) based on likelihood and Markov chain Monte Carlo (MCMC). Also in this approach, the model has an analytically intractable likelihood function meaning that the probability of the observed data as a function of the parameters of interest is not available in closed form. The reason for the intractability is the presence of unobserved variables (forming the transmission tree in her model), which are integrated out using MCMC. While feasible for a small number of variables, this technique runs into problems when the number of unobserved variables is large (see e.g. Green *et al*. 2015). The problems manifest themselves in the form of increased convergence issues of the Markov chain. Aandahl *et al*. (2014) resolved such issues in the approach of Stadler (2011) but concluded that the ABC method has a similar accuracy and better computational efficiency than the amended version.

In all of the tests comparing distance measures in ABC, the generalized Kullback-Leibler distance measure attained lower or equal estimation error compared to the baseline measure introduced by Tanaka *et al*. (2006), suggesting that one can reduce the estimation error to some degree by the choice of the distance measure only (see also Fearnhead and Prangle 2012). A possible disadvantage of the approach with the generalized Kullback-Leibler distance is a slight increase in computational cost. Although not an issue in our study where the computational bottleneck was the simulator, it may be an issue when the inference procedure is repeated for a large number of data sets, or when the simulation of the data is not computationally expensive. While the measure used by Tanaka *et al*. (2006) has some clear biological meaning, the generalized Kullback-Leibler distance is based on a more general information-theoretical construction. The observed increase in the performance is thus interesting, because in ABC, distances are usually strongly based on application-specific knowledge even though some exceptions do exist (Gutmann *et al*. 2014).

Our explicit construction of the approximate likelihood function put on display the difficulties in the estimation of *R* when inferring both the reproductive value *R* and the transmission rate *t* (Tanaka *et al*. 2006; Blum 2010). The credible intervals for *R* were (1.4, 38.0) or (9.5, 80.0) when using a uniform prior over the (*α, δ*) or (*R, t*) space, respectively. The large upper end points of the credible intervals reflect the extreme flatness of the approximate likelihood function with respect to the reproductive value parameter. An uninformative prior over the (*α, δ*) space has been, to our knowledge, the standard choice in all of the related ABC studies. Comparing the posterior with the approximate likelihood function shows how significantly the prior contributed to the posterior, altering the shape of the likelihood function greatly. The results highlight the usefulness of likelihood approximation as an identifiability check when performing inference for models with intractable likelihoods.

A uniform prior is usually considered uninformative so that it may seem paradoxical that the prior had such a strong influence on the posterior. The apparent paradox is readily resolved by noting that the uniform prior was not imposed on the actual epidemiological quantities of interest but on a nonlinear transformation of them.

The standard assumption in the previous ABC studies concerned with the Tuberculosis data from San Francisco (Small *et al*. 1994) has been that the infectious population size *m* equals 10,000 individuals. We showed that the infectious population size influences the estimation of *R*, with the estimate of *R* increasing with *m*. Assuming *m* = 10, 000, the posterior mean was 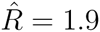. Smaller assumed population sizes *m* = 1000 and *m* = 5000 yielded smaller estimates 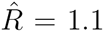 and 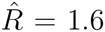, respectively, while larger assumed population size *m* = 20, 000 increased the estimate to 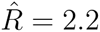. The corresponding likelihoods (proportional to posterior distributions with uniform prior) with varying *m* were clearly distinct from each other as visible in Figure 3.

Taking advantage of the dependency between *R* and *m*, we showed that it is possible to estimate the infectious population size *m* when *R* and *δ* are known. Using the estimate *δ* = 0.52 (Aandahl *et al*. 2014) and assuming either *R* = 2.1 or *R* = 1.1, the posterior means of *m* were 21, 100 or 2100, respectively, for the San Francisco data of Small *et al*. (1994). Further biological expertise can be used to assess the reasonability of different inferred population sizes in a comparable modeling setting.

We noticed that for small values of *m*, the generative model was unable to produce data with a similar number of distinct alleles as the San Francisco data while containing also large clusters, supporting the observation of Tanaka *et al*. (2006) that *m* = 1000 does not result in an appropriate level of diversity. In the San Francisco data, large clusters were present originating from groups of people with conditions affecting the immune system, e.g. AIDS. Among such groups the transmission rate of *M. tuberculosis* can be expected to be notably higher and thus rapidly producing large clusters. The simple model, however, does not account for these situations. In future work it would be interesting to consider approximate inference for models with possibly heterogeneous reproductive values that depend on auxiliary epidemiological data (Bacaër *et al*. 2008). However, given the apparent identifiability issues with the simple model studied here, it would be of utmost importance to ensure that the molecular and epidemiological data are jointly informative enough to perform reliable inferences.

## ACKNOWLEDGMENTS

This research was funded by the Academy of Finland (Finnish Centre of Excellence in Computational Inference Research COIN). We acknowledge the computational resources provided by the Aalto Science-IT project.

## APPENDIX A

Stabilization of the approximate likelihoods:

## APPENDIX B

Mean and median absolute errors:

The rather large difference between the mean and median errors of *R* in Figure 6 indicates that there are some large errors which pull the mean error up. This is mostly due the tendency of *R* = *α/δ* to be large when the estimate of the death rate *δ* is small. Although the errors for *R* in Figure 6 are large, the small errors in Figure 7 indicate that the estimation of the transmission rate *t* can still be done accurately and is not affected by the error-prone estimate of *R*.

**Figure 6:**
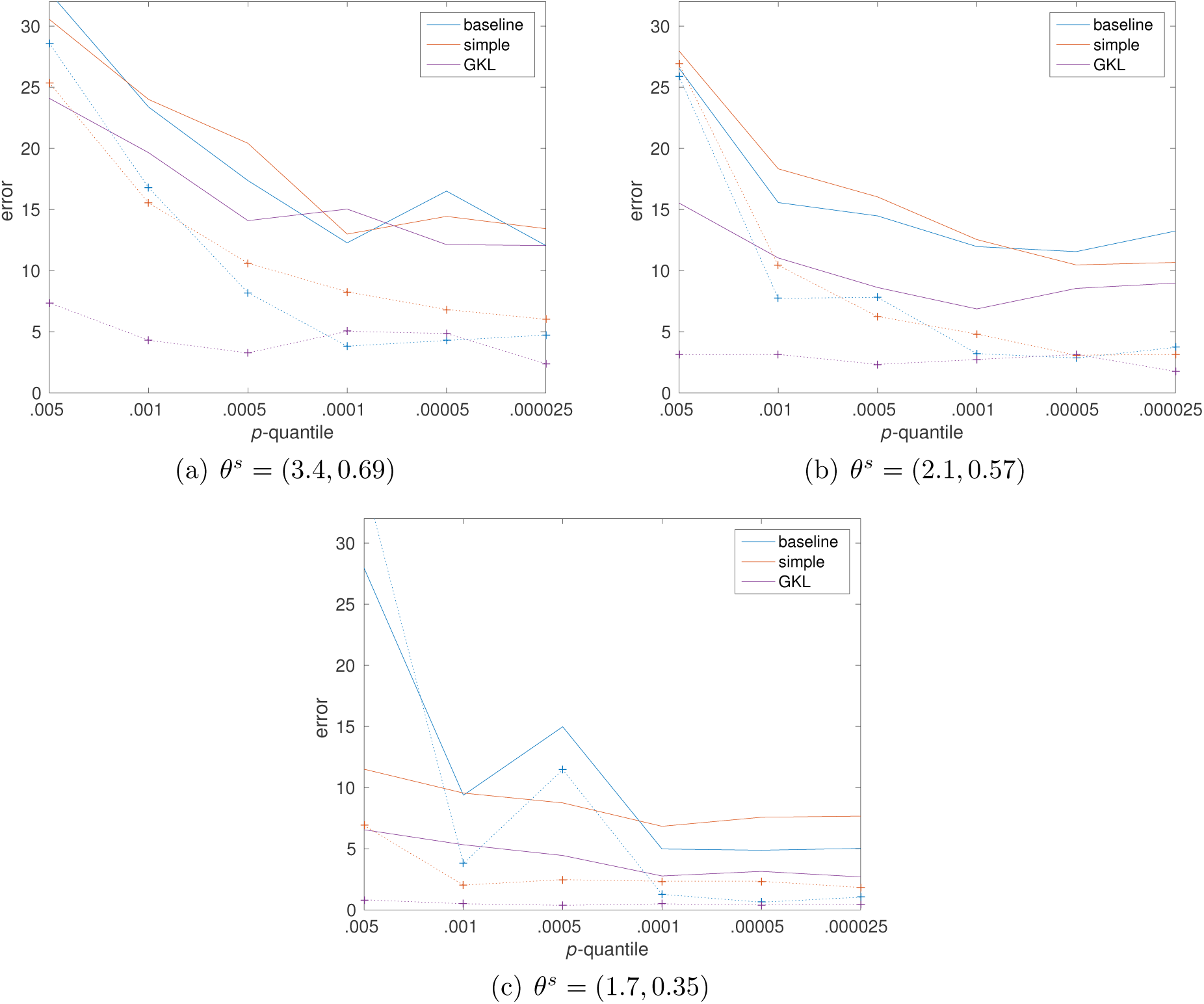
Mean (solid lines) and median (dotted lines) errors of the approximate maximum likelihood estimates with the three different distance measures d_base_ (blue lines), d_sim_ (red lines), d^GKL^ (purple lines) and a decreasing threshold *ϵ* given by the *p*-quantile. The error is the *L*_1_ distance between *R^s^* and 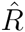, that is, 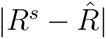.

**Figure 7:**
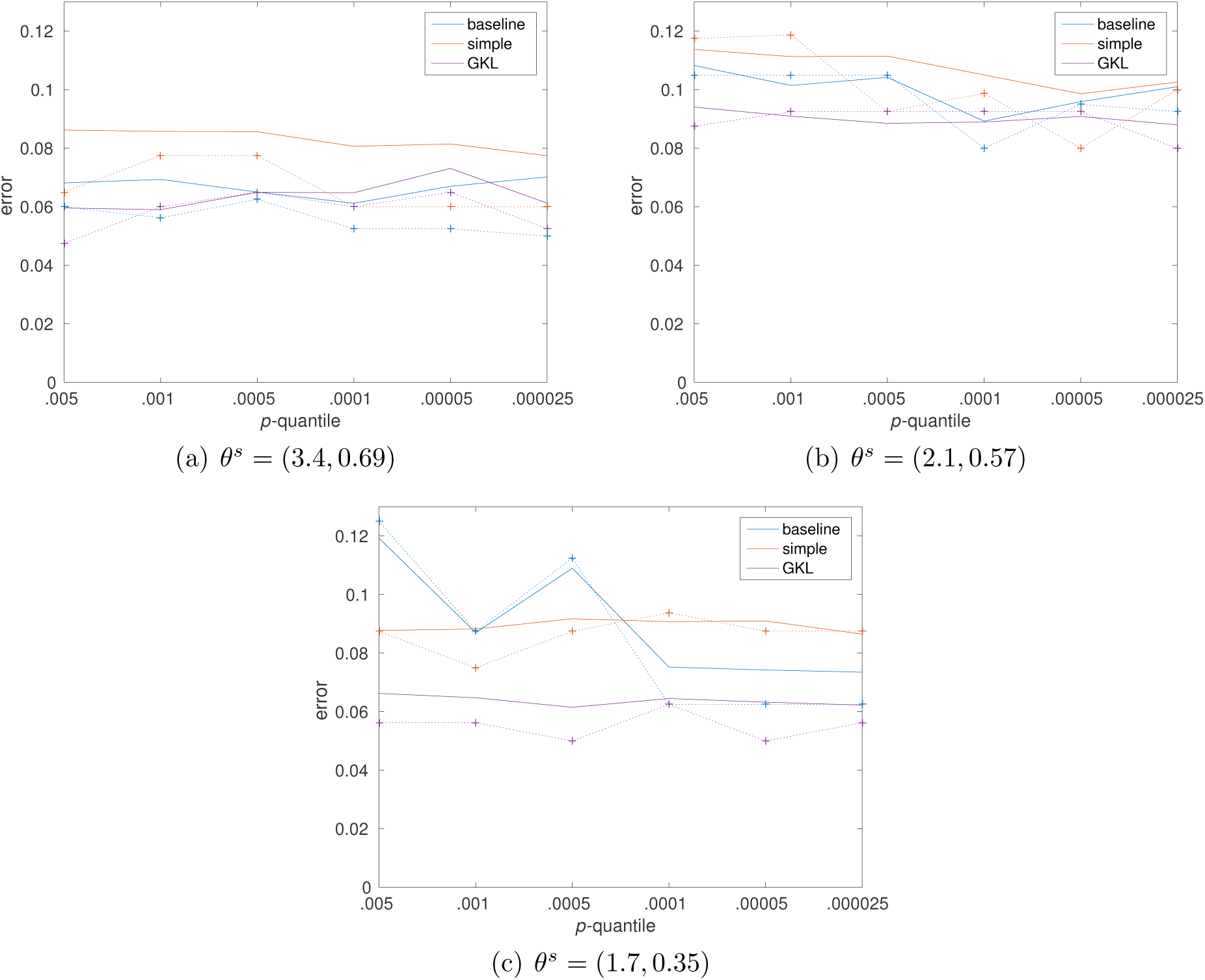
Mean and median errors of the approximate maximum likelihood estimate of *t*. Visualization is as in Figure 6. The error is the *L*_1_ distance between *t*^s^ and 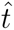, that is, 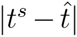

Absolute errors in the rate parameter space:

We noticed a general tendency of acquiring small estimates of *δ* irrespective of the setup. This is a plausible reason for the slightly reduced errors in Figure 8 (a) compared to the other setups as *δ*^*^ is the smallest in that setup.

**Figure 8:**
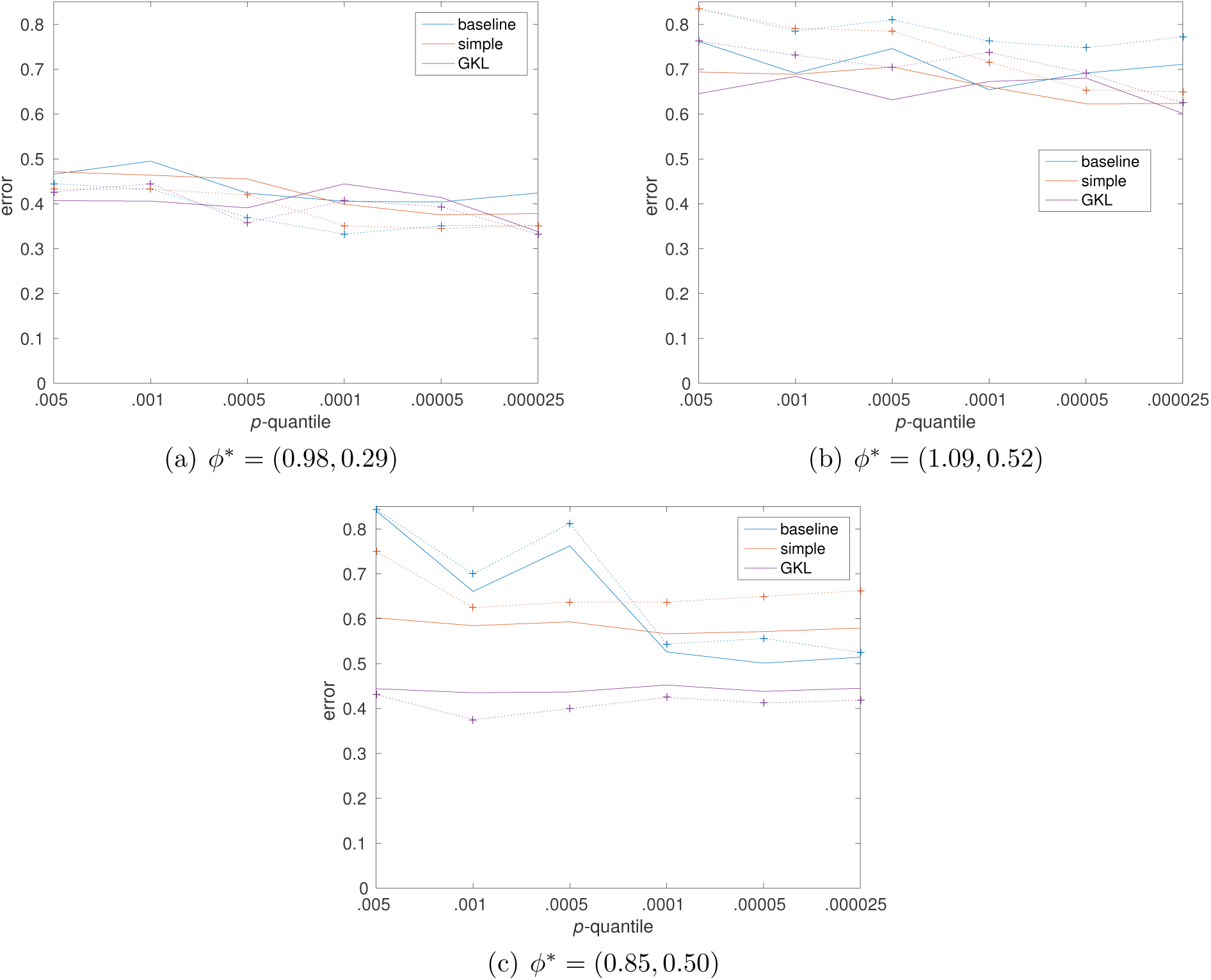
Mean and median errors in the estimated *ϕ* = (*α, δ*). Visualization and setup is as in Figure 6. The error is the *L*_1_ distance between the vectors *ϕ*\* and 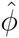.

## APPENDIX C

Transformed uniform prior:

**Figure 9:**
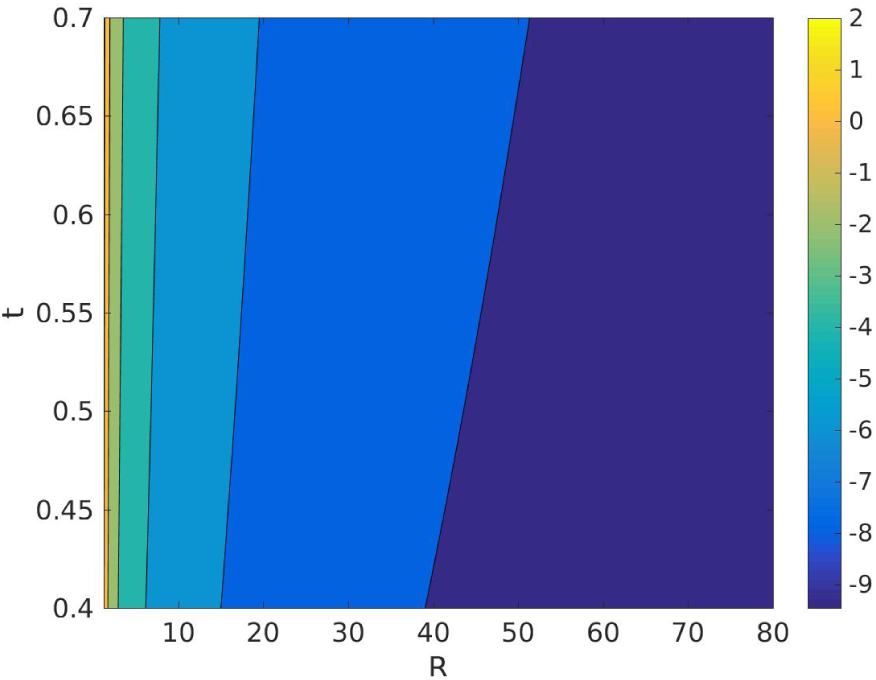
Visualization of the logarithm of the prior in Equation 5 corresponding to the uniform prior on (*α, δ*). Note the concentration of probability mass on small *R*.

